# Visualization and Morphological Analysis of Individual Oligodendrocytes in the Mouse White Matter

**DOI:** 10.1101/2025.10.27.684952

**Authors:** Yasuyuki Osanai, Sasikarn Looprasertkul, Batpurev Battulga, Reiji Yamazaki, Nobuhiko Ohno

## Abstract

Myelin formation by oligodendrocytes is essential for the regulation of the conduction velocity and proper brain function. To ensure accurate information processing in response to experiences such as sensory stimuli and learning, oligodendrocytes adjust their number and morphology. In addition, oligodendrocyte morphology changes with senescence and in the presence of neurodegenerative diseases. Thus, visualizing oligodendrocytes and analyzing their morphology is crucial for understanding how our brains change under such conditions. Herein, we describe the methods for labeling and analyzing the morphologies of individual oligodendrocytes in mouse white matter at the light microscopic level. PDGFRa-CreERT2:Tau-mGFP and PLP-CreERT2:Tau-mGFP mice enable us to visualize and analyze later-born or early-born oligodendrocyte morphology. In addition, sparse oligodendrocyte labeling with attenuated rabies virus expressing GFP enables the visualization and morphological analysis of individual oligodendrocytes in various brain white matter regions without the need for transgenic animals. Furthermore, the combination with immunostaining in thick tissues enables the identification of labeled oligodendrocytes and myelin sheaths, as well as their interactions with neuronal axons. These methods are suitable for revealing how oligodendrocytes adapt their morphologies depending on environmental stimuli or pathological conditions.

## 1. Introduction

Oligodendrocytes, the myelin sheath-forming glial cells in the central nervous system, are critical for proper brain functions as they regulate the conduction velocity through myelin sheath formation and saltatory conduction and support neuronal metabolism. Physiological and pathological changes in brain functions are often accompanied by the structural regulation of oligodendrocytes and their myelin, which in turn modulates signal transduction in the neural circuits. Neuronal axons can be divided into multiple segments, including internode covered by myelin sheaths and nodes of Ranvier. The nodes of Ranvier are flanked by paranodes involving paranodal loops and axon-oligodendrocyte junctions [1]. Internodes are characterized by multilamellar compact myelin and inner and outer tongue cytoplasmic channels. Each oligodendrocyte, typically with a small cell body, extends multiple processes, with each connected to the outer tongue [2]. Characteristic oligodendrocyte morphology is developed through the differentiation of oligodendrocyte progenitor cells (OPCs), which express markers such as NG2 and PDGFRa and exhibit bipolar shapes, into mature myelinating oligodendrocytes, which test positive for markers such as proteolipid protein (PLP) and myelin basic protein, extend multiple processes, and form myelin sheaths.

Compared to other glial cells such as astrocytes and microglia, visualization of oligodendrocytes is difficult because there are no reliable anti-bodies that label both their cell bodies and processes (including myelin). Historically, silver carbonate methods or single-cell injection of lucifer yellow have been used for visualizing the morphology of a single oligodendrocyte [3, 4]. However, as silver carbonate staining requires special fixation and single-cell lucifer yellow injection is technically difficult and makes it hard to obtain high sample numbers, recent studies have focused on the use of transgenic animals and viral vectors [5–7].

PDGFRa-CreERT2:Tau-mGFP and PLP-CreERT:Tau-mGFP mice are transgenic lines that label oligodendrocytes differentiated after tamoxifen administration or those generated before tamoxifen administration, respectively [8, 9]. The PDGFRa-CreERT2 mouse expresses tamoxifen-inducible Cre in OPCs, and the Tau-mGFP mouse possess the lox-STOP-lox membrane-targeted GFP (mGFP) cassette downstream of the Tau (MAPT) promoter that is active in neurons and mature oligodendrocytes. Tamoxifen-induced recombination removes the lox-STOP-lox cassette from OPCs, and as they differentiate into oligodendrocytes, GFP expression is driven by the Tau promoter. Thus, in PDGFRa-CreERT2:Tau-mGFP mice, we can label the oligodendrocytes differentiated after the tamoxifen injection. In contrast, in PLP-CreERT mice, tamoxifen-inducible Cre is expressed only in mature oligodendrocytes, since the PLP promoter is active in mature oligodendrocytes but not in OPCs. Therefore, PLP-CreERT:Tau-mGFP mice enable the labeling of oligodendrocytes differentiated before the tamoxifen injection. Thus, in PLP-CreERT:Tau-mGFP mice, we can label early-born oligodendrocytes when we administer tamoxifen in the early developmental period [9]. Therefore, these transgenic animals enable the visualization of a temporarily distinct oligodendrocyte population.

Rabies virus and Semliki Forest virus vectors have been used to sparsely label oligodendrocytes in the mouse white matter [10, 11]. We have previously analyzed the interactions between oligodendrocytes and other cells, such as neurons, using a rabies virus vector and immunostaining for thick (> 70µm) sections. Using this method, both single oligodendrocytes and their myelinated axons can be labeled [9]. Thus, it is a powerful tool to analyze oligodendrocyte-neuron interactions.

In this paper, we introduce methods to visualize and analyze single-cell oligodendrocyte morphology at the light microscopic level, focusing on two labeling methods employing transgenic mice and a viral vector. The methods using PDGFRa-CreERT2:Tau-mGFP and PLP-CreERT:Tau-mGFP have been validated in the corpus callosum and cerebellum [9, 12], and labeling using the rabies virus vector has been employed in multiple white matter regions, including the corpus callosum, cerebellum, optic nerve and chiasm, and spinal cord [10]. Methods utilizing these transgenic animals are potentially applicable to all brain regions, including the gray matter. The rabies virus-mediated method is considered applicable not only to the white matter of mice but also to that of other animals. These methods can be useful in examining various morphological aspects of individual oligodendrocytes, such as the number of myelin sheaths, the length and fiber diameter of each myelin sheath, height of each oligodendrocyte, and myelination toward a particular subtype of axons.

## 2. Materials

### 2.1 Transgenic mice and reagents

1. PDGFRa-CreERT2 [13] or NG2-CreERT2 mice (Jackson, strain name: B6.Cg-Tg(Cspg4-cre/Esr1*)BAkik/J, # 008538) [14].
2. PLP-CreERT mice (Jackson, strain name: B6.Cg-Tg(Plp1-cre/ERT)3Pop/J, # 005975) [15].
3. Tau-mGFP mice (Jackson, strain name: B6;129P2-Mapttm2Arbr/J, # 021162) [16].
4. Tamoxifen.
5. Corn oil.
6. Disposable syringes for small-volume injection.
7. 25G short length needles (10–16 mm).

### 2.2 Viral vector and stereotaxic injection

1. Rabies B19G-SADΔG-GFP (Salk Institute, viral vector core) or Rabies B19G-HepΔG-GFP [17] (*See* Note 1 for information on virus production and purchasing).
2. Apparatus for stereotaxic microinjection (For detailed procedures of mouse brain stereotaxic injection, please consult methodological papers on viral injection, such as Cetin et al., 2006) [18].

### 2.3 Perfusion and slicing of the brain

1. 4% paraformaldehyde (PFA) in 0.1 M phosphate buffer (PB) (pH 7.4).
2. Vibratome slicer.
3. Perfusion pump.

### 2.4 Thick tissue immunostaining and tissue clearing with RapiClear

Stock solutions without normal goat serum (NGS) or antibodies can be stored at room temperature (RT; 20–25°C) for over 1 year.

1. Penetration buffer: 2% Triton X-100 and 0.05% sodium azide in phosphate-buffered saline (PBS). Mix 2 g of Triton X-100 and 50 mg of sodium azide in 98 mL of PBS.
2. Blocking buffer: 10% NGS, 1% Triton X-100, 2.5% dimethyl sulfoxide (DMSO), and 0.1% sodium azide in PBS. Mix 1 g of Triton X-100, and 2.5 g of DMSO in 87 mL of PBS. Add NGS at one-tenth of the final volume to the solution before use.
3. Antibody dilution buffer: 1% NGS, 0.2% Triton X-100, 2.5% DMSO and 0.1% sodium azide in PBS. Mix 2 mL of 10% Triton X-100 solution, 2.5 g DMSO, and 100 mg sodium azide in 96 mL of PBS. Add NGS at 1/100 of the final volume of the solution immediately before use.
4. Washing buffer: 3% NaCl and 0.2% Triton X-100 in PBS. Mix 3 g of NaCl, and 2 mL of 10% Triton X-100 solution in 95 mL of PBS.
5. Antibodies: Mouse monoclonal anti-APC (CC1, 1/100, IgG2b, OP80, Merck) for labeling mature oligodendrocytes, rabbit polyclonal anti-human Olig2 (1/100, 18953, Immuno-Biological Laboratories America) or chicken polyclonal anti-Olig2 (1/500, OLIG2-0020, Aves) for labeling oligodendrocyte lineage cells, rabbit polyclonal anti-Iba1 (1/400, 019-19741, Wako) for labeling microglia, mouse monoclonal anti-GFAP (1/100, IgG1, G3893, Sigma-Aldrich) or mouse polyclonal anti-S100B (1/100, ab34686, Abcam) for labeling astrocytes (*see* Note 2), rabbit polyclonal anti-Caspr (1/500-1000, ab34151, Abcam) for labeling the paranodes of the myelin sheath, rabbit polyclonal anti-NG2 (1/50, AB5320, Merck) for labeling OPCs, rabbit polyclonal anti-calbindin (1/500, MSFR100390, Frontier Institute) for labeling Purkinje cells, chicken polyclonal anti-neurofilament-H (1/500, NB300-217, Novusbio) for labeling neuronal axons, and rat monoclonal anti-GFP (1/500, IgG2a, 04404-84, Nacalai Tesuque) for visualizing GFP.
6. RapiClear 1.52 (RC152001, SUNJin Lab, Taiwan)
7. Antifadant

## 3. Methods

### 3.1 Labeling oligodendrocytes using PDGFRa-CreERT2:Tau-mGFP and PLP-Cre:Tau-mGFP mice

#### 3.1.1 Tamoxifen administration and fixation

1. PDGFRa-CreERT2 and PLP-CreERT mice are crossed with Tau-mGFP mice to generate PDGFRa-CreERT2 (either homozygous or heterozygous):Tau-mGFP (heterozygous) and PLP-CreERT (either homozygous or heterozygous):Tau-mGFP (heterozygous) mice, respectively.
2. Tamoxifen is dissolved in corn oil for one 1 h to make a tamoxifen stock solution (10 mg/mL). During this process, the solution is incubated at 60 ℃ and vortexed every 10 min. Tamoxifen stock solution can be aliquoted (0.5–1.5 mL) and stored for at least six months at −20℃. The frozen stock solution is incubated at 60℃ for 1 h before use, with gentle mixing by vortex every 10 min during incubation. The thawed solution can be stored at 4℃ for up to one week. Before injection, the solution is incubated at 60℃ for 10 min and vortexed thoroughly. Ideally, the temperature of the solution be 37℃ at the time of injection.
3. Tamoxifen is administered via intraperitoneal injection (50–100mg/kg) for two consecutive days in mice younger than postnatal day 15 (P15). For mice older than P15, intraperitoneal administration of 100 mg/kg tamoxifen for four consecutive days is recommended (*see* Notes 3, 4, and 5).
4. Mice are perfused with 4% PFA in 0.1 M PB at the desired postnatal age. Their brains are then postfixed in the same fixative for 1 day at 4℃. After transferring the brains into PBS, 100 µm-thick brain slices are prepared using a vibratome slicer (*see* Note 6).

### 3.2 Labeling oligodendrocytes using attenuated rabies virus-GFP

The methods for stereotaxic microinjections are well established; therefore, they are not described in detail here. For the detailed procedure, please refer to methodological papers on stereotaxic injection, such as Cetin et al (2006).

1. Dilute the rabies virus stock with PBS to prepare a viral solution containing 5.0 × 10^3^–5.0 × 10 ^4^ infectious units per microliter.
2. Inject 1 µL of the viral solution into the white matter. The following parameters are used for injections into each brain region. In the adult mouse, for the corpus callosum 1.0 mm posterior and 0.5–0.8 mm lateral to the bregma, at a depth of 1.0 mm from the pia, for the optic chiasm, 0.2 mm posterior and 0.0 mm lateral to the bregma, at a depth of 5.0–5.3 mm from the pia, and for the white matter of cerebellar lobules IV/V, 6.45 mm posterior and 1.00 mm lateral to the bregma, a depth of 0.5 mm.
3. Perfuse mice with 4% PFA in 0.1 M PB, 4–7 days after viral injection. The brains are then postfixed in the same fixative for 1 day at 4℃. After transferring the brains into PBS, 100 µm-thick brain slices are prepared using a vibratome slicer.

### 3.3 Immunostaining of thick brain sections using the Rapi clear method to analyze oligodendrocyte morphology

1. Select brain sections (50–200 µm) containing the target brain region and place 2–3 sections per well into a 24-well plate filled with PBS.
2. Incubate brain sections in the penetration buffer at RT for 24 h.
3. Incubate the brain sections in the blocking buffer for 1–2 days at 4℃,
4. The sections are incubated at 4℃ for 3–4 days in the antibody dilution buffer (300-500 µL for 2–3 sections in 24 well plate) containing primary antibodies such as rat anti-GFP (1/500), rabbit anti-Caspr (1/500-1000), rabbit anti-human Olig2 (1/100) or chicken anti-Olig2 (1/500), and mouse anti-APC (CC1, 1/100).
5. Wash the sections in washing buffer three times for 1 h each at RT. Then, incubate the specimens in washing buffer on an orbital shaker at 4℃ overnight.
6. Incubate the specimens with secondary antibodies in the antibody dilution buffer on an orbital shaker at 4℃ for 2 days.
7. Wash the sections in washing buffer three times for 1 h each at RT. Then, keep the specimens in the washing buffer on an orbital shaker at 4℃ overnight.
8. Stain with Hoechst (1/1500–2000) in PBS for 3 h at RT if needed.
9. If the tissue was stained with Hoechst, wash it with PBS three times for 1 h each on an orbital shaker at RT. If not, with PBS twice for 10 min each.
10. For tissue clearing, place the specimen on a slide glass, add pre-warmed (37℃) RapiClear solution with/without an antifade solution. After 20 min, applying the cover glass.
11. Apply the cover glass and seal its edges with a clear nail polish. Allow the slide to dry overnight at RT (*see* Note 7).

### 3.4. Confocal microscopic imaging

Use a confocal microscope with Z-series sequential imaging function. A single oligodendrocyte morphology can be captured by approximately 50–100 optical sections at 0.5 µm intervals, adjusted as needed to fully capture the dorsal-ventral extent of the cell bodies, processes, and myelin sheaths of individual oligodendrocytes. We utilized a 60× objective lens (N.A. 1.42) on the Dragonfly 200 high-speed confocal platform to achieve the high– magnification image. High–resolution images are required for detailed morphological analyses of individual oligodendrocytes and their myelin internodes.

1. The uppermost and lowermost individual oligodendrocytes and their myelin are set for image acquisition, with a step size of 0.5 µm. GFP signal loss is used to define the uppermost and lowermost limits of the confocal Z-stack imaging.
2. Images are captured and exported in the proprietary file format provided by the microscope manufacturer. The subsequent analysis is performed using the Fiji software, which supports most proprietary formats, such as .oif, .nd2, .czi, and .ims. These formats preserve z-plane and scale information, facilitating the analysis of entire individual oligodendrocytes with Fiji.

### 3.5. Analysis of oligodendrocyte morphology labeled using PDGFRa-CreERT2:Tau-mGFP and PLP-Cre:Tau-mGFP mice

The oligodendrocyte in PDGFRa-CreERT2:Tau-mGFP (Fig. 1) and PLP-Cre:Tau-mGFP mice (Fig. 2) are labeled via tamoxifen-induced Cre recombination at specific time points. The oligodendrocyte cell membrane and myelin sheath membrane can be labeled by mGFP, which enables the analysis of the oligodendrocyte’s morphology. The age of mice, number of administrations, and concentration of the tamoxifen are important for the analysis of a single oligodendrocyte. If oligodendrocytes are densely labeled, it becomes difficult to analyze the morphology of individual cells. Various morphological aspects can be analyzed in single oligodendrocytes (*see* Note 8), such as the myelin sheath length, number of myelin sheaths produced by a single oligodendrocyte [8, 19], fiber diameter of each myelin sheath [20], and heights of single oligodendrocytes [12].

**Fig 1.**
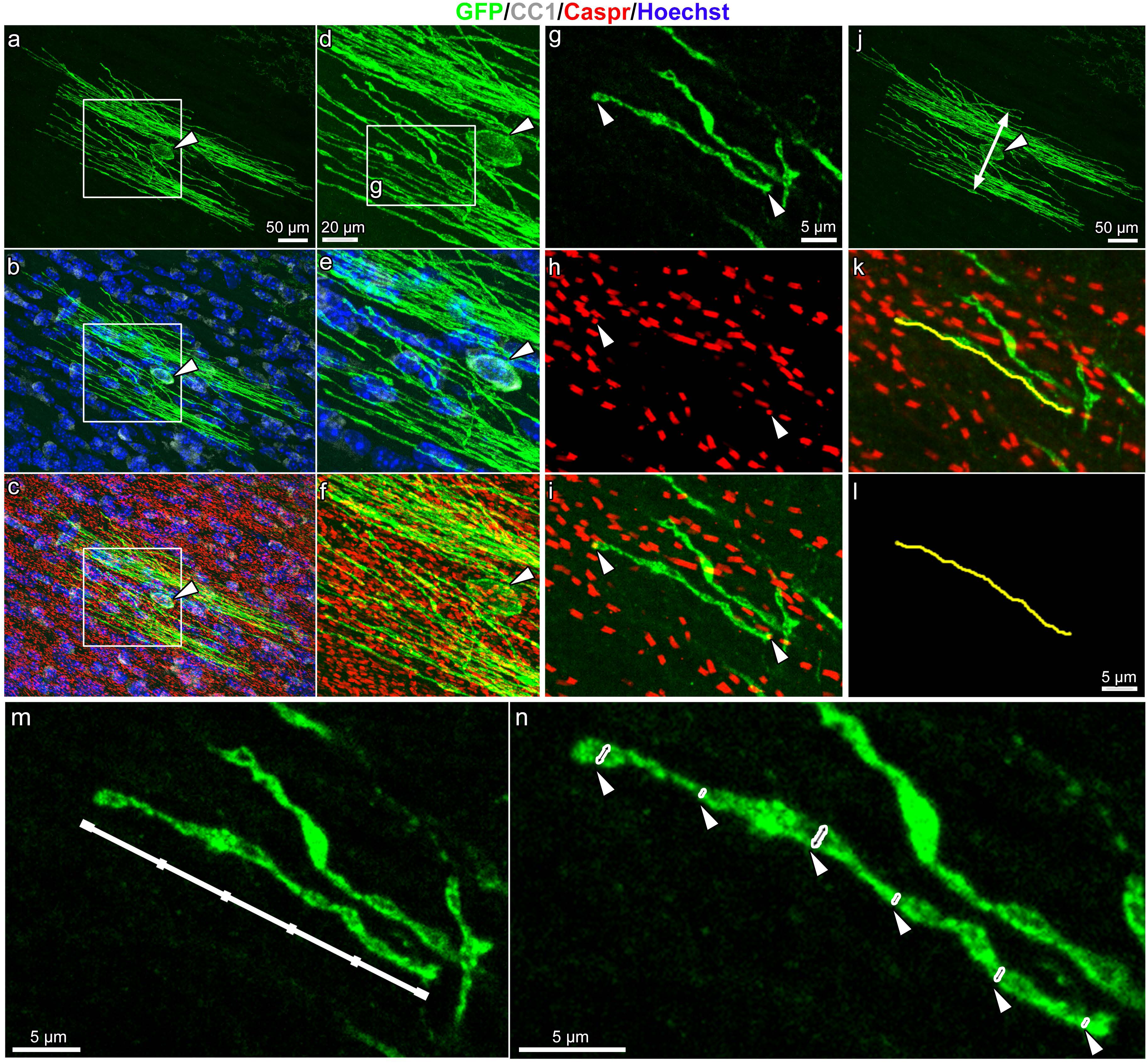
Visualization and morphological analysis of individual oligodendrocytes in the corpus callosum of PDGFRa-CreERT2:Tau-mGFP double transgenic mice. (a-f) Representative z-stacked images of an oligodendrocyte in the corpus callosum of 10-week-old PDGFRa-CreERT2:Tau-mGFP mice that were injected tamoxifen two weeks before perfusion. The tissue was immunostained for GFP (green), CC1 (white), Caspr (red), counterstained with Hoechst (blue), and imaged using a confocal microscope. The rectangles in the left panels (a–c) are magnified in the right panels (d–f). (g–i) A higher-magnification z-stacked image of the area marked by the rectangle in d. The image shows a GFP-labeled myelin sheath (green, g and i) flanked by Caspr-positive paranodes (red, arrowheads, h and i). This method illustrates a single myelin sheath observable in the z-stacked serial images. (j) GFP-labeled oligodendrocyte (green, arrowhead) in a z-stacked image; double-headed arrow indicates cell height. (k–l) A z-stacked image shown in i with the yellow line added in the analysis with the SNT plugin in Fiji. The GFP-positive myelin sheath (green, k), flanked by Caspr-positive paranodes (red, k), is traced with a yellow line (k and l), representing the path used to measure the length of a single internode formed by an individual oligodendrocyte. This tracing was performed across z-stacked serial images. (m–n) Representative images showing fiber diameter measurement (axonal diameter + myelin thickness) of a single internode shown in g. A GFP-labeled myelin sheath of the individual oligodendrocyte was selected and traced along its full length using a z-stacked serial image. A segmented line was then drawn on a single plane of the z-stacked image and divided into five segments, starting 1 µm from the tip of each side (the white line and its segment lines, m). Fiber diameters were measured at six points along each internode (the arrowheads indicate measurement sites and double-headed arrows indicate span of the fiber diameter, n).

**Fig 2.**
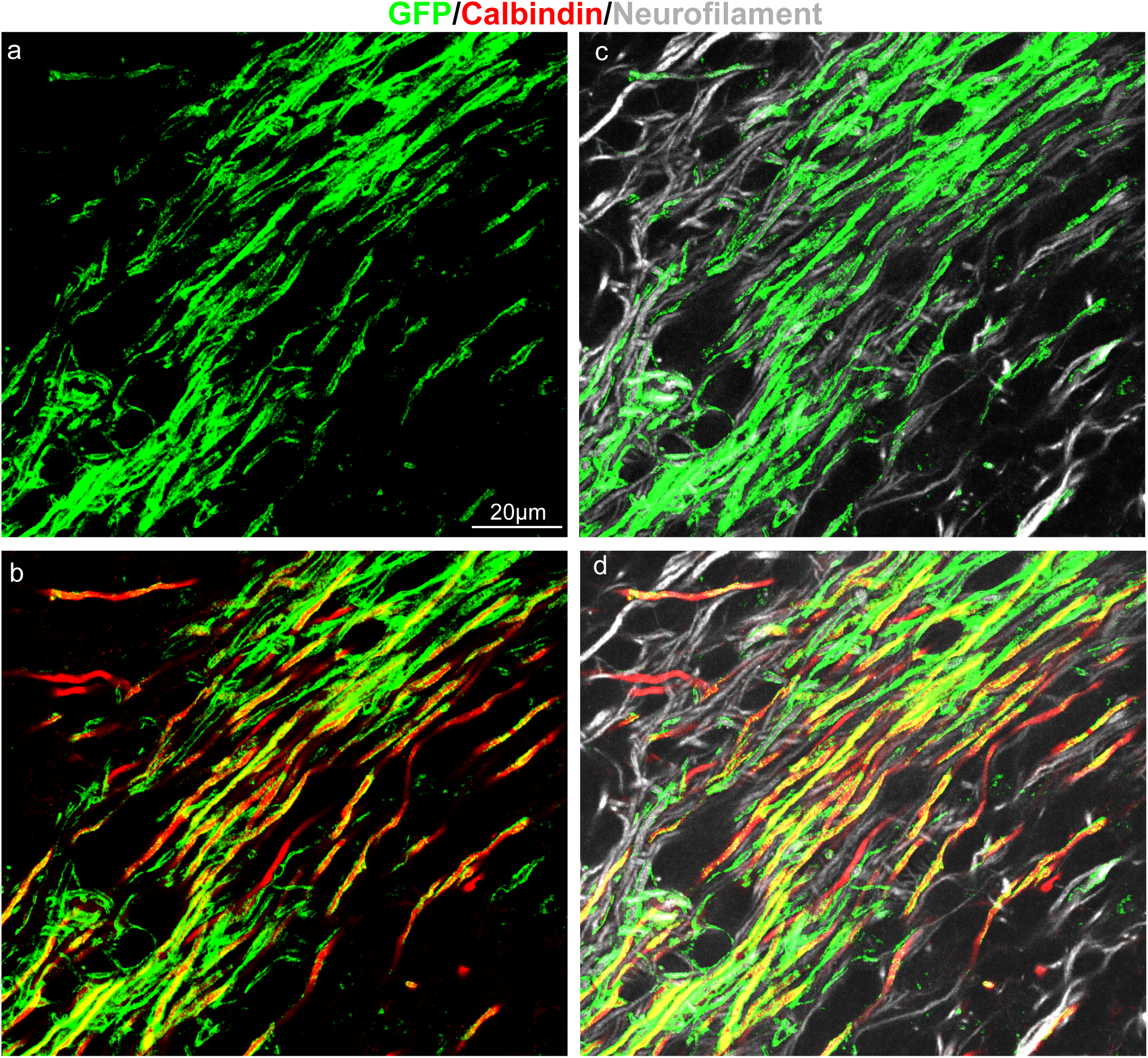
Analysis of Purkinje cell selective myelination by early-born oligodendrocytes using PLP-CreERT:Tau-mGFP double transgenic mouse. Representative single-plane confocal image of cerebellar white matter of an 8-week-old PLP-CreERT:Tau-mGFP double transgenic mouse treated with a single tamoxifen injection at P8. (a) GFP-labeled early-born oligodendrocytes and their myelin sheaths (green). (b) Merged image showing Purkinje cell axons (calbindin, red) and early-born oligodendrocytes (GFP, green). (c) Neuronal axons (neurofilament, white), including Purkinje cell axons and other axon subtypes, along with early-born oligodendrocytes (green). (d) Confocal image showing Purkinje cell axons (red), neurofilament-positive axons (white), and early-born oligodendrocytes with their myelin sheaths (green). Due to close proximity of labeled oligodendrocytes, individual cells cannot be clearly distinguished. However, since individual myelin sheath can be identified, we enable to assess whether early-born oligodendrocytes preferentially myelinate Purkinje cell axons (*see* note 5).

#### 3.5.1 Measurement of the length and number of myelin sheaths formed by a single oligodendrocyte

1. The number of sheaths and the myelin sheath length produced by a single oligodendrocyte are quantified using the Simple Neurite Tracer (SNT) plugin (https://imagej.net/plugins/snt/) [21], accessed via Fiji via Plugins > Neuroanatomy > SNT….
2. In the SNT Startup Prompt window, select the directory containing the target file that includes the z-stacked serial images of the GFP-labeled oligodendrocyte and its myelin sheaths.
3. Since the images may appear with low intensity, adjust the brightness and contrast as needed via Fiji > Image > Adjust > Brightness/Contrast….
4. Each individual myelin sheath is manually traced in the z-stacked serial images. The starting point of each myelin sheath is defined at one tip of the myelin sheath (GFP-positive) colocalized with the paranode marker (Caspr-positive, Fig. 1g–i), and subsequent points are selected along the trajectory of the sheath across adjacent optical sections. Press either “Yes” or the “Y” key to accept the path. For each pair of selected points, the A* search path-finding algorithm auto-tracing within SNT is used to compute the optimal path based on image intensity gradients. Each path segment is accepted by pressing the “Y” key. After completing a full trace of one sheath, the path is finalized by selecting “Finish” or pressing the “F” key.
5. Each completed myelin sheath tracing is stored as an individual path and can be verified using SNT Path Manager (Fig. 1k, l).
6. After tracing all the myelin sheaths of a single oligodendrocyte, save the tracing data as follows: File > Save Tracings… > Save As… (.traces file).
7. To determine the length of each myelin sheath and the total number of sheaths per cell, export the path statistics via Analysis > Path Properties: Export CSV….

#### 3.5.2 Height/ vertical territory measurement of an individual oligodendrocyte

1. Acquire z-stacked serial images of individual oligodendrocytes at a 0.5 µm step size, ensuring that all myelin sheaths of the cell are included. Images should be saved in a file format compatible with the Fiji software.
2. Import the z-stacked serial images into the Fiji software. The imported images display the entire oligodendrocyte, with all its myelin sheaths.
3. Since the images may appear with low intensity, adjust the brightness and contrast appropriate via Fiji > Image > Adjust > Brightness/Contrast….
4. Identify the outermost dorsal and ventral myelin processes of the oligodendrocyte using the z-stack images.
5. Locate the uppermost dorsal and lowermost ventral myelin sheaths by tracing along their full length and identifying the connection point between each myelin sheath and the corresponding oligodendrocyte process.
6. Mark these connection points for further analysis.
7. Locate and mark the nucleus of each individual oligodendrocyte in the z-stack images.
8. Compress the confocal z-stack into a single-plane projection for linear distance measurement via Image > Stacks > Z Project…
9. Measure the distance between the points where the outermost dorsal and ventral myelin internodes connect to their process by drawing a straight line from the dorsal outermost myelin sheath to the ventral outermost one. Ensure that the line passes through the nucleus and is perpendicular to the connection points of the processes (Fig. 1j). This can be done via Analyze > Measure.

#### 3.5.3 Fiber diameter measurement of a single oligodendrocyte

1. Acquire z-stacked serial images of individual oligodendrocytes at a 0.5 µm step size, ensuring the inclusion of all myelin sheaths of the cell. Images should be saved in a file format compatible with the Fiji software.
2. Import the z-stacked serial images into the Fiji software. The imported images display the entire oligodendrocyte, including all its myelin sheaths.
3. Since the images may appear with low intensity, adjust the brightness and contrast appropriate via Fiji > Image > Adjust > Brightness/Contrast….
4. Select the myelin sheaths of each oligodendrocyte to measure their diameters. One approach is to include: (i) the outermost myelin sheath on the dorsal side, (ii) the outermost sheath on the ventral side, (iii) one internode located in the central region of the oligodendrocyte, (iv) one internode between the central region and the outermost dorsal sheath, and (v) one internode between the central region and the outermost ventral sheath, based on z-stack images.
5. Compress the Z-stacked image to a single plane image. To determine where to measure the diameter, draw a line parallel to the myelin internode of the selected myelin sheath and measure the length of that line (Fig. 1m).
6. Since the fiber diameters can vary along the internode, myelin internodes are divided into several segments for measurement. For example, the internodes are divided into six segments using the following formula to determine the spacing along their total length: Distance for measurement position (µm) = (Internode length (µm) − 2 µm) / 5 Place the first and last measurement points at a distance of 1 µm from each end of the myelin internode because the internode’s tip is a paranode area that is always larger than the internode. The remaining four points are positioned equally between them based on the calculated distance.
7. After the positions for the measurement are marked (Fig. 1m), draw a line from the upper part to the lower part of that myelin area and measure its length (Fig. 1n) via Analyze > Measure.
8. Collect data on the selected positions of each myelin sheath and calculate the average fiber diameter.
9. Measure the fiber diameter of the selected internodes per oligodendrocyte—approximately five internodes per oligodendrocyte. Then, calculate the average fiber diameter of each oligodendrocyte.

### 3.6 Analysis of selective myelination toward a particular subtype of axons

We have previously demonstrated that oligodendrocytes that differentiated early in life preferentially myelinate Purkinje cell axons in the cerebellar white matter [9]. By combining the RapiClear method with oligodendrocyte labeling using transgenic mice or rabies virus, we are able to investigate the selective myelination toward specific axon subtypes.

1. Label oligodendrocytes using transgenic mice or rabies virus. We labeled early-born oligodendrocytes in PLP-CreERT:Tau-mGFP mice by administering tamoxifen at P8 (100 mg/kg) and perfusing the animals with 4% PFA in 0.1 M PB at P56 (Fig. 2).
2. Prepare 100 µm brain slices using a vibratome.
3. Performing thick-section immunostaining, RapiClear method, as described above, using markers for specific neuronal subtypes (e.g., calbindin for Purkinje cell axons) and the pan-neuronal axon marker neurofilament. GFP-labeled oligodendrocytes in transgenic animals should be enhanced with an anti-GFP antibody, whereas those labeled by rabies virus do not require immunostaining.
4. Acquire z-series confocal images of GFP-labeled oligodendrocyte/myelin and multiple subtypes of neuronal axons as described above.
5. Count the number of axons of each subtype myelinated by the GFP-labeled sheath using Fiji. GFP-labeled processes ensheathing axons for more than 20 µm were defined as myelin sheaths (*see* note 9).
6. Count the number of axons of each subtype not myelinated by GFP-labeled oligodendrocyte/myelin within the height of the labeled oligodendrocytes (Fig. 2).
7. To determine whether GFP-labeled oligodendrocytes preferentially myelinate specific axon subtypes, perform Fisher’s exact test for individual confocal images using statistical software (e.g., Prism, GraphPad). Input the following values:
  A. Number of specific-subtype axons myelinated by GFP-labeled oligodendrocytes,
  B. Number of specific-subtype axons not myelinated,
  C. Number of non-specific-subtype axons myelinated,
  D. Number of non-specific-subtype axons not myelinated.

Fisher’s exact test is then performed to evaluate selective myelination.

### 3.7 Analysis of single oligodendrocytes morphology labeled by attenuated rabies virus

While the morphological analyses of individual oligodendrocytes in the sections labeled with the rabies virus are mostly similar to those in PDGFRa-CreERT2:Tau-mGFP and PLP-CreERT:Tau-mGFP mice, there are some differences. The GFP expression level in cells infected by attenuated rabies virus encoding GFP (hereafter referred to as RV-GFP) is so strong that immunostaining for GFP is unnecessary. Since RV-GFP can label cells other than oligodendrocyte-lineage cells, we first need to check whether the cells labeled with RV-GFP are oligodendrocytes. If the labeled cells exhibit morphological features of myelinating mature oligodendrocytes and have GFP-positive ensheathment around neuronal axons, which are mostly flanked by Caspr immunostaining, these cells are identified as oligodendrocytes.

In the white matter, myelinating mature oligodendrocytes labeled with RV-GFP extend the processes corresponding to their myelin sheaths parallel to the direction of the neuronal axons. Because RV-GFP labels the cytosol of myelin sheaths, the ensheathment of the GFP signal around unstained axons can be clearly observed when the diameters of these axons are large (Fig. 3a, *see* Note 9). Another distinguishing feature of RV-GFP-labeled oligodendrocytes is the swelling observed at the paranodal regions (Fig. 3). Since the paranodes contain more cytoplasm than the internodes, the paranodes appear more intensely labeled and expanded with GFP. The length of each myelin sheath can be measured by identifying its paranodes as GFP-labeled swellings at both ends of the sheath.

**Fig 3.**
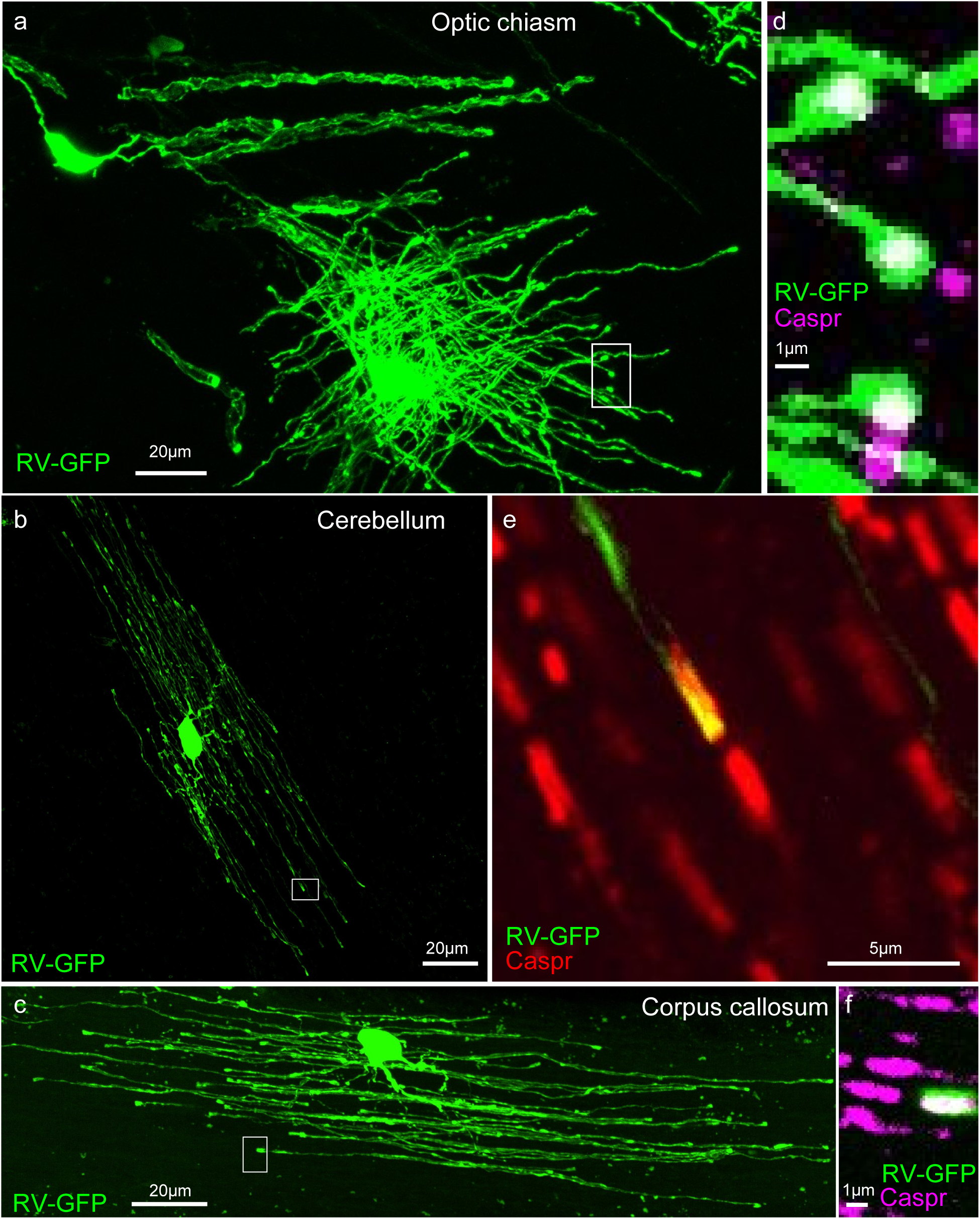
Oligodendrocytes labeled by RV-GFP. (a–c) RV-GFP-labelled oligodendrocytes in the optic chiasm. (a), cerebellar white matter (b), and corpus callosum (c) of 8-week-old mice perfused 4 days after injection. (d–f). Double-fluorescence images of the GFP and Caspr signals in the region outlined by the white square in panels a–c. respectively. The distal ends of processes are co-localized with Caspr signals, indicating that the processes are myelin sheaths.

We can also use antibodies against oligodendrocytes/myelin markers. Notably, cell body markers for oligodendrocytes (e.g., CC1) tend to be downregulated in RV-GFP (+) cells, which might be attributed to the high-level GFP expression. To confirm whether GFP-labeled cells form the myelin sheath, we recommend using an anti-Caspr antibody. If the ends of GFP-positive processes are closely aligned with Caspr signals, this indicates that the processes correspond to the myelin sheath flanked by the set of Caspr signals (Fig. 3). These morphological features, combined with immunostaining, allow for reliable oligodendrocyte identification.

## 4. Notes

1. Salk Institute viral vector core provides a stock G-deleted rabies aliquot. Rabies virus vector amplification techniques have been described in several studies [22, 23]. Rabies B19G-SADΔG-GFP and rabies B19G-HepΔG viral vectors can be obtained from Vector Builder (Chicago, USA).
2. Immunostaining of membrane-located receptors is not always successful for the thick-tissue immunostaining method. For example, immunostaining for PDGFRa is difficult with this method. If you want to count the number of OPCs, we recommend quantifying CC1(−)/Olig2 cells [24] or using immunostaining for NG2 [12]. In our experience, the anti-GFAP antibody exhibits reliable performance in brain tissue, whereas the anti-S100B antibody lacks astrocyte specificity, particularly in the corpus callosum. In contrast, in the optic nerve, anti-GFAP antibodies are ineffective, but the anti-S100B antibody works well. To accurately determine the number of astrocytes in the optic nerve, double immunostaining with anti-S100B and anti-Olig2 antibodies is recommended, as both astrocytes and OPCs express S100B.
3. In our experience, 100 mg/kg tamoxifen administration for more than two days increases the risk of mortality in mice younger than P15. Researchers should first confirm whether the mouse strain they are using can tolerate two consecutive days of tamoxifen administration. If the mortality rate is high, the number of tamoxifen administration days or its dosage should be reduced.
4. In our experience, tamoxifen administration before P9 failed to label oligodendrocytes; instead it labeled unidentified cells in PDGFRa-CreERT2:Tau-mGFP mice, possibly because PDGFRa promoter activity is not yet restricted to OPC at such an early developmental stage.
5. Four days of consecutive injection of tamoxifen at 100 mg/kg helps maximize the recombination rate. If a lower recombination rate is acceptable or if you wish to limit the number of labeled oligodendrocytes, consider reducing the tamoxifen concentration or the number of injection days. In PLP-CreERT:Tau-mGFP mice, oligodendrocytes in the white matter tend to be densely labeled [9]. Therefore, to achieve sparse labeling and enable observation of single-cell morphology, the tamoxifen concentration must be substantially reduced.
6. Brain slices can be stored for up to six months at 4℃ in PBS containing 0.1% sodium azide or stored for up to two years at −80℃ after freezing in PBS containing 30% sucrose. Before freezing, the slices are incubated in PBS containing 30% sucrose overnight. The GFP signal of Tau-mGFP mouse tissue is weak and needs to be enhanced by immunostaining for GFP prior to imaging.
7. The immunostained sections embedded on slides can be stored at 4℃ for up to one month. For longer storage, the slides need to be stored at −20℃.
8. During the tracing process, it is important to distinguish the GFP-positive myelin sheaths of the target oligodendrocyte from those of neighboring cells. Because adjacent oligodendrocytes can also express GFP, their myelin sheaths may appear to be in close proximity, potentially leading to misidentification. To avoid this issue, for the morphological analysis of single oligodendrocytes, only cells that are isolated within the field of view (regions where a single GFP-positive oligodendrocyte was clearly identifiable) are selected for tracing. Furthermore, only myelin sheaths directly connected to the cell body through proximal processes of the targeted oligodendrocyte are included in the analysis.
9. Myelination analyses performed on large-caliber axons, such as cerebellar white matter and the optic nerve, are easier and more reliable compared to those performed on small-caliber axons, such as the axons in the corpus callosum, because we can clearly observe GFP-positive processes ensheathing neuronal axons (Figs. 2 and 3) [25].

